# A derived ZW chromosome system in *Amborella trichopoda*, the sister species to all other extant flowering plants

**DOI:** 10.1101/2020.12.21.423833

**Authors:** Jos Käfer, Adam Bewick, Amélie Andres-Robin, Garance Lapetoule, Alex Harkess, José Caïus, Bruno Fogliani, Gildas Gâteblé, Paula Ralph, Claude W. dePamphilis, Franck Picard, Charlie P. Scutt, James Leebens-Mack, Gabriel AB Marais

## Abstract

Sex determination is poorly understood in plants. *Amborella trichopoda* is a well-known plant model for evo-devo studies, which is also dioecious (has male and female individuals), with an unknown sex determination mechanism. *A. trichopoda* is a “sex switcher”, which points to possible environmental factors that act on sex, but populations grown from seed under greenhouse conditions exhibit a 50:50 sex ratio, which indicates the operation of genetic factors. Here, we use a new method (*SDpop*) to identify sex-linked genes from genotyping data of male and female individuals sampled in the field, and find that *A. trichopoda* has a ZW sex-chromosome system. The sex-linked genes map to a 4 Mb sex-determining region on chromosome 9. The low extent of ZW divergence suggests these sex chromosomes are of recent origin, which is consistent with dioecy being derived character in the *A. trichopoda* lineage. Our work has uncovered clearly formed sex chromosomes in a species in which both genetic and environmental factors can influence sex.

**One Sentence Summary:** *Amborella trichopoda*, a dioecious species in which both genetics and the environment influence sex, possesses a pair of quite recently evolved ZW chromosomes.

## Main Text

Dioecy (separate male and female individuals) is one of the many sexual systems exhibited by flowering plants. This system has evolved repeatedly in angiosperms, and about 6% of angiosperm species are dioecious (*1, 2*). Genetic sex determination (GSD) has been reported in about 50 angiosperm species (*3, 4*) and master sex-determining genes have been identified in a few of these (*5–9*). Despite these great advances, we still lack a global understanding of the evolution of sex determination in dioecious plants. In particular, it is not clear how widespread GSD is, nor whether there are common genetic dynamics contributing to the evolution of dioecy in flowering plants. Environmental sex determination (ESD), defined as the possibility of a plant to switch sex during its lifetime (*10, 11*), has been poorly studied, probably because it is thought to be very rare (*1, 12*). However, even in species with well-established sex chromosomes, sex switching and incomplete sex determination are not uncommon (*2, 13*–*15*). It has been speculated that ESD is a consequence of gene expression changes elicited by environmental triggering of phytohormone signaling pathways (*11*), but we simply have no data on ESD, nor on the mechanisms responsible for environmentally induced floral phenotype changes in plants with GSD systems.

*Amborella trichopoda*, a dioecious shrub endemic to New Caledonia, is the sole species on the sister lineage to all other angiosperms, and as such has been used in evo-devo investigations of floral evolution (*16*–*18*). Rigorous ancestral state reconstruction suggests that the last common ancestor of all extant angiosperms had bisexual flowers (*19*) and independent transitions to dioecy have occurred repeatedly throughout angiosperm evolution (*1*), including within the *Amborella* lineage. Despite broad interest in *A. trichopoda* given its pivotal phylogenetic position, the nature and timing of the evolutionary shift to dioecy on the lineage leading to *Amborella* has remained enigmatic. Sex-switching is well documented in *A. trichopoda* (*20–22*) suggesting some degree of ESD. Whereas sex ratios in wild populations are often male-biased, plants grown from seeds exhibit a sex ratio close to 1:1 upon first flowering, suggesting a combination of GSD and ESD (*22*). No evidence for sex chromosomes has been observed in the *A. trichopoda* karyotype, but many plant sex chromosomes are homomorphic, i.e. not distinguishable in light microscopy (*3*). Here we combine a novel approach to identify sex-linked markers in natural populations with genetic mapping to test hypothesized GSD and characterize the *A. trichopoda* sex chromosomes.

Identifying sex chromosomes from genomic data can be difficult, especially when these are homomorphic (*4, 23*). First, we investigated the presence of sex-linked markers in a sample of 10 female and 10 male individuals grown from seeds collected in the wild. RNA-seq data of these individuals were obtained from flower buds and used to genotype them. We used a novel method, *SDpop*, which uses modifications of the Hardy-Weinberg equilibrium expectation to identify sex-linkage (*24*). This allows probabilistic classification of genes as autosomal or sex-linked with *XY* or *ZW* heterogametic genotypes. The data obtained in this study strongly supported the *ZW* model (Table 1), inferring 49 *ZW* gametologs and 9 *Z*-hemizygous genes (lacking a detectable *W* copy).

**Table 1.** Results of the *SDpop* analysis of the RNA-seq data from a natural population

|  | # genes<br>with<br>expression<sup>*</sup> | # genes<br>with<br>informativ<br>e SNPs<sup>□</sup> | #SNPs | BIC<br>without sex<br>chromosom<br>es | BIC XY | BIC ZW | #<br>autoso<br>mal<sup>§</sup> | # z-<br>hemi<br>zygo<br>us<sup>§</sup> | #<br>ZW<sup>§</sup> |
| --- | --- | --- | --- | --- | --- | --- | --- | --- | --- |
| **unmasked** | 24460 | 17223 | 420101 | 5599123 | 5594811 | 5590632 | 7518 | 9 | 49 |
| **NoW** | 24415 | 17195 | 420766 | 5601968 | 5597448 | 5590920 | 7741 | 9 | 62 |
| **NoZ** | 24418 | 17206 | 420047 | 5443303 | 5438903 | 5432196 | 7696 | 9 | 57 |

\* at least 1 read
□ at least 1 individual of each sex had more than three reads, and the gene had at least 1 SNP
§ based on the ZW model, posterior probability threshold for classification is 0.8, BIC = Bayes Information Criterion

We tested the *SDpop* inference by mapping putatively sex-linked genes to a chromosomal draft assembly of the *A. trichopoda* genome (V.6, genome_ID = 50948, https://genomevolution.org/coge) including 13 pseudomolecules corresponding to each chromosome pair (n=13). We mapped the 49 gametologous genes identified by *SDpop* onto this genome and found that 33 (67%) of them mapped uniquely to a single region of ∼4 Mb on chromosome 9 of the assembly (Figure 1). This proportion increased to 89% when genomic contigs not included in the 13 chromosome scaffolds were excluded from the reference.

**Fig. 1.**
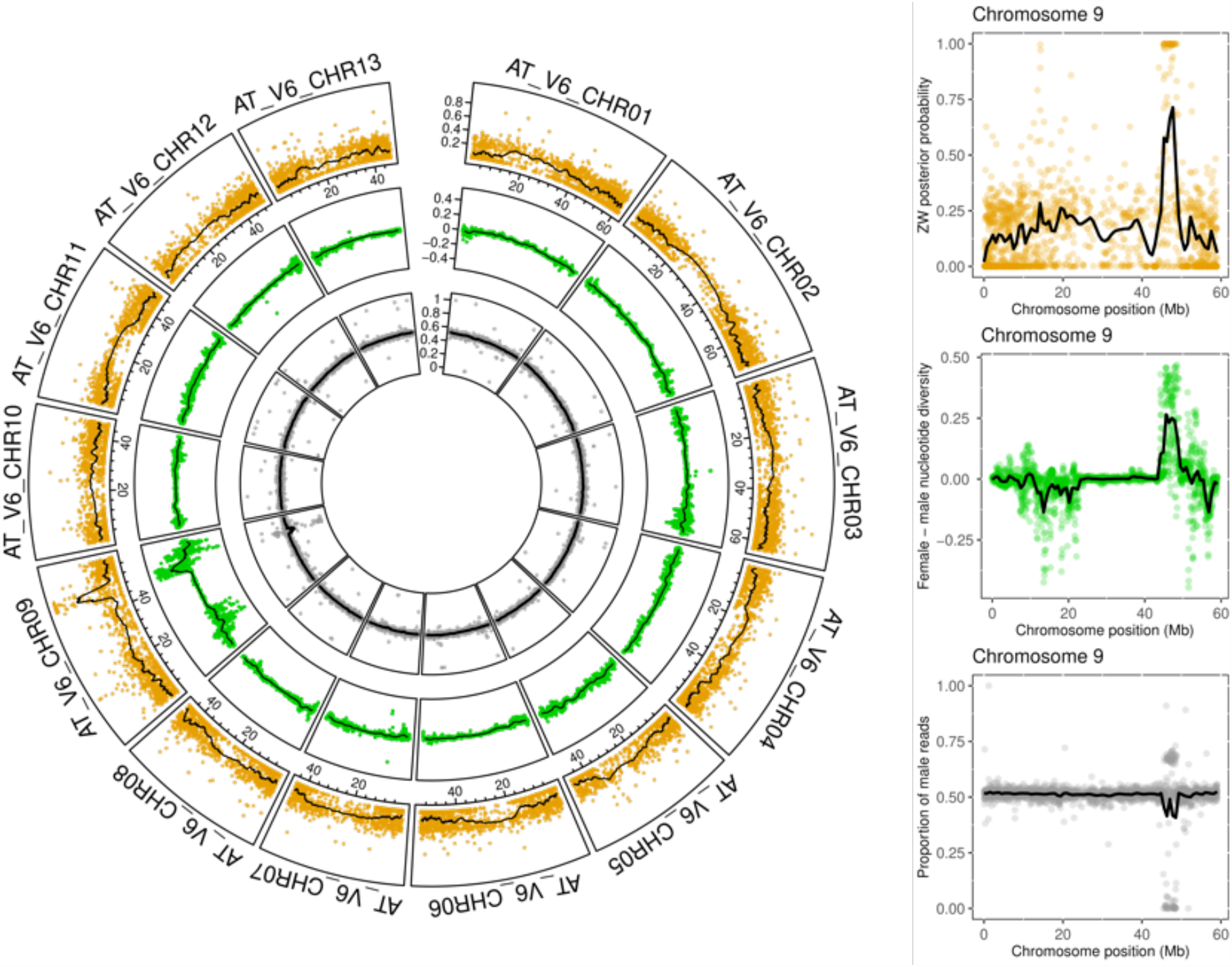
Evidence for a ZW sex-determining region in *A. trichopoda*. (A) Circular representation of the *A. trichopoda* genome. Chromosome number (1 to 13) is indicated. Outer circle indicates ZW posterior probability, inner circle indicates female – male nucleotide diversity, in-between circle indicates proportion of male reads. (B) Zoom in for chromosome 9. The sex-determining region is located at positions 45-49 Mb (see text for more details). Region 10-20 Mb shows an excess of male heterozygous – female homozygous sites reminiscent of a XY locus, which explains a drop in female – male nucleotide diversity. This region is in fact not associated to sex and is discussed in Supplementary Text.

We investigated the extent of the sex-determining region further by mapping re-seq data from a male, a female and 33 F1 progeny for which flower sex had been scored (16 female and 17 male F1 offspring). First, we assessed whether 671 female-specific SNP variants identified in the wild population transcripts were only present in the seed parent and female progeny in the mapping population. Males in the mapping population were found to possess no more than five of the putatively female-specific SNP variants identified in the wild-collected population, whereas females were found to possess at least 465 of these variants (Table S1).

We also used the 33 genotyped and phenotyped progeny to estimate the frequency of the sites that were heterozygous in females and homozygous in males, and found a clear excess of female-heterozygous SNPs in the sex-determining region identified by *SDpop* (Figure 1). Finally, the male-versus-female coverage ratio(*4, 23*) was found to deviate significantly from one in the 4 Mb sex-determination region identified by *SDpop* (Figure 1).

*SDpop* and heterozygosity analyses performed on the wild-sampled plants and mapping population, together with an assessment of the mapping population read coverage data, provided consistently strong support for an approximately 4 Mb-long region on the right arm of chromosome 9 (45-49 Mb) as cosegregating perfectly with sex. However, we found another region, on the left arm of chromosome 9 (10-20 Mb), showing a weaker association with sex (Figure 1). The recombination landscape on chromosome 9 may explain this observation (Supplementary Text, Figure S1).

The individual that was sequenced in the *Amborella* genome project (*SantaCruz 1975*.*3*) is known to have switched from producing mostly male flowers (*20*) to all female flowers (*18*). The mapping of reads from the male and female progeny array described above identified the presence of both *Z* and *W* gametologs in the clearly *ZW*, genetically female reference genome. Further scrutiny of the sex-determining region revealed that the reference assembly of this region is chimeric with a mixture of Z and W segments. The male/female coverage-ratio analysis mentioned above was used to identify Z and W segments and construct draft Z and W haplotypes by masking W-linked scaffolds (with only female reads, and thus a male/female coverage-ratio ∼ 0) and Z-linked scaffolds (with a male/female coverage-ratio ∼ 2), respectively (Supplementary Text, Figure S2). Remapping the reads on these masked assemblies reduced the regions with sex-biased coverage (DNA-reseq data) and yielded more sex-linked genes (Table 1).

The sex-determining region includes ∼150 sex-linked genes and functional analyses will be required to determine how many of these may influence floral phenotypes. The ability of *A. trichopoda* genotypes (including *SantaCruz 1975*.*3*) to produce both male and female flowers or change sex completely (*20*–*22*) suggests that both the Z and W regions include genes that singly or collectively can act as a switch to induce male or female flower developmental programs. We focused on i) ZW gene pairs with differential expression in males and females, and ii) genes present in one haplotype and not the other (W-specific or Z-hemizygous genes). We used the RNA-seq data from flower buds mentioned above to conduct a male vs female differential expression analysis (Figure S3). Although a few genes in the sex-determining region had sex-biased expression, none were expressed exclusively in males or females (Data S1). We identified 37 genes exhibiting sex-biased expression. We then looked at the annotation of these genes (Data S2) and found that four of them were involved in the auxin pathway, a known sex-related phytohormone (*25*) (Table S2). The fact that all four genes are male-biased suggest a possible masculinizing effect. We also looked at the putative W-specific and Z-hemizygous genes that we identified (Data S1) and found that a W-specific gene had a DMY domain that is characteristic of PPR genes (Data S2), a class of genes involved in cytonuclear interactions, and particularly in cytoplasmic male sterility (*26*) (Table S2).

We used masked assemblies to calculate the synonymous divergence (*dS*) between the two copies of all predicted genes in the assembly and then used the maximum Z-W *dS* values obtained to estimate the age of the sex-determining region (*4, 27*). The highest *dS* values are estimated from the mapping population re-seq data, and are around 0.14, but these values have large standard errors. The values estimated from the natural population RNA-seq have lower maximum values and standard errors. Among-gene variation in *dS* is high, as expected due to sampling variation (*28*). We therefore estimate that the maximum *dS* value is between 0.04 and 0.06 (Figure 2). This is 20 to 30 times higher than the level of genome-wide nucleotide polymorphism (0.002), suggesting that recombination suppression occurred 9.5 to 14.5 times 4\**Ne* generations ago. Pairwise sequentially Markovian coalescent (PSMC) (*29*) analysis of SNP variation among extant *A. trichopoda* populations yield a *Ne* approximation of 37,500 and past population size fluctuations (*18*). When using the extant *Ne* estimate and a generation time of four years (*18*), we estimate suppression of recombination in the *A. trichopoda* sex-determination region of the *Amborella* chromosome 5.7 to 8.7 million years ago. Four years is probably a lower bound estimate of the *A. trichopoda’s* generation time in the wild as individuals of 40-50 years old have been observed (*30*). A much more realistic estimate is probably around 20-25 years and the sex-determining region could be as old as 54.3 millions years. The divergence of the *Amborella* lineage from that of all remaining extant angiosperm is estimated to have occurred at least 125 million years ago (*31, 32*), and the mean *dS* value between *A. tricophoda* and other angiosperms was estimated to be between 1 and 2.2 (*33*). We found a much smaller *dS* for the sex-determining region, which points to a much more recent origin than the split of the *Amborella* lineage from the other angiosperms.

**Fig. 2.**
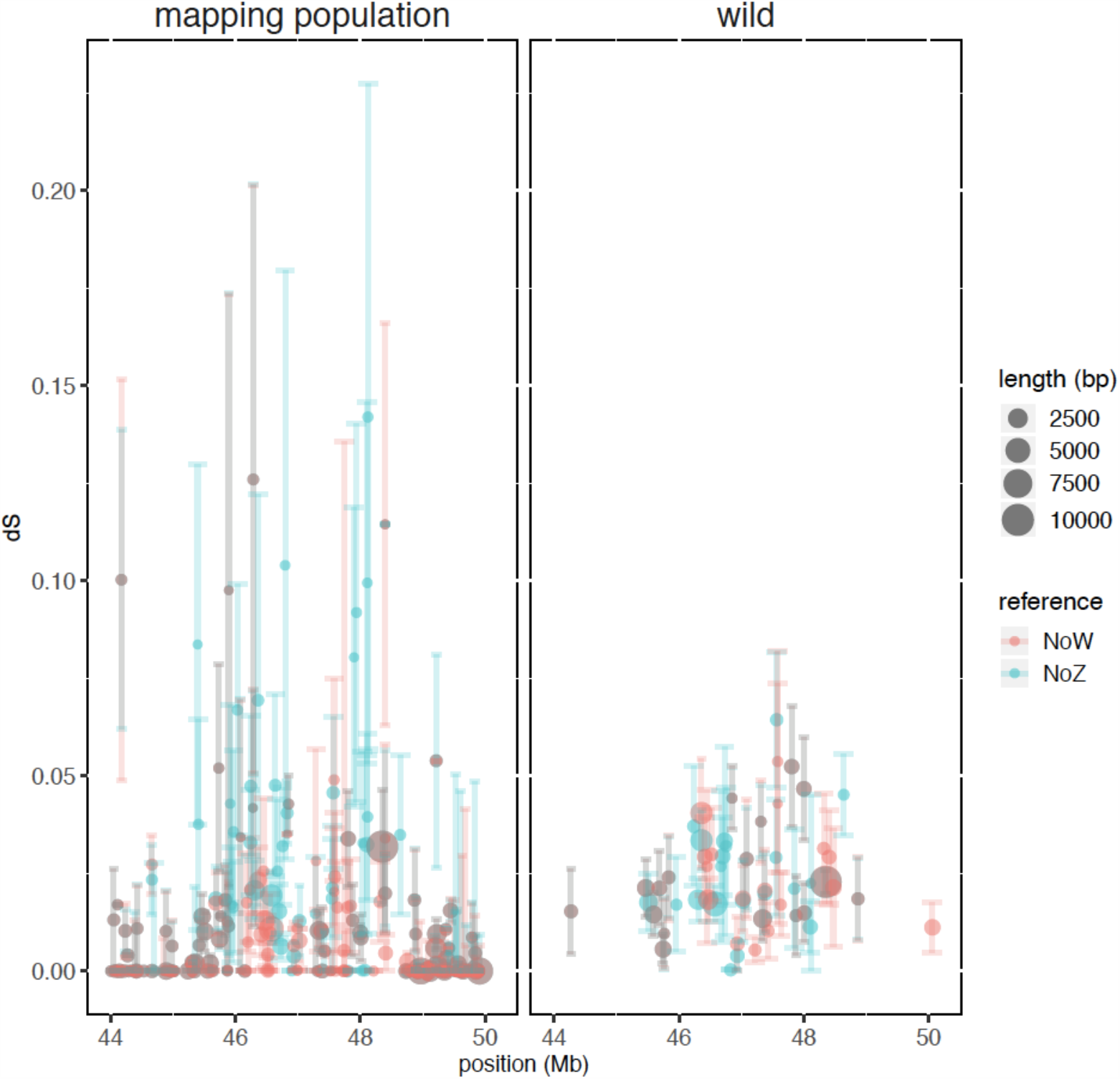
Synonymous divergence between Z and W gametologs along the sex-determining region. A) in the mapping population and B) in the collection of plants from a wild population. The synonymous divergence is the *dS* values obtained using codeml on coding sequences (see Material and Methods). Coding sequence length is indicated. Data are shown using both assemblies (Z masked and W masked, see text for more details).

Our results clearly point to the existence of a *ZW* system with a small (∼4 Mb) sex-determining region on chromosome 9 of *A. trichopoda*. Thus, *Amborella* has genetic sex determination, and all of the 55 individuals used in this study had a phenotypic sex that corresponded with their genetic sex. However, as in many plant species with sex chromosomes, sex changes can occur, as illustrated by the individual used for the reference genome and observations described in previous studies (*22*). The reasons why the plant used for the reference genome did not initially correspond to its genetic sex, and why it subsequently switched to its genetic sex, are both unknown. It is, however, known that local heating of branches of an otherwise female *Amborella* plant can induce male flowering in these branches; thus, genetic sex determination doesn’t rule out any influence from the environment on sex. A previous study suggested that *A. trichopoda* has inconstant males, which could indicate that dioecy had evolved from a gynodioecious intermediate in this species (*22,34*). However, the observation of sex-switching and the influence of the environment on sex could indicate that genetic sex determination evolved from environmentally controlled monoecy, in which plant hormones might have played an important role (*25*). The presence of distinct Z and W haplotypes in *Amborella* could be used to test how often phenotypic and genetic sex differ in nature, and whether inconstant individuals always have the same genetic sex.

*A. trichopoda* is the sole extant species in the sister lineage to all other extant flowering plants. This species and other members of the ANA grade (including Nymphaeales and Austrobaileyales) are of pivotal importance for reconstructing evolution of flower development and the characteristics of the ancestor of all angiosperms (*16,18*). Based on extent angiosperm species, it was inferred that the most likely ancestral breeding system was bisexuality (*19*). However, several ANA-grade species such as some *Trithuria* spp. (Nymphaeales) and *Schisandra chinensis* (Austrobaileyales) are dioecious (*35,36*). Dating the origin of dioecy in these plants is very important to accurately reconstruct the ancestral sexual system (*22*). Our estimate of the age of the ZW chromosomes of *A. trichopoda* points to relatively recent sex chromosomes compared to the chronology of angiosperm evolution, which does not support the view that dioecy is ancient in *A. trichopoda*. However, we cannot completely rule out this possibility, as frequent sex chromosome turnover is possible in some lineages (*37,38*). The structure of the *A. trichopoda* flower, however, does not fit well with ancient dioecy, as non-functional male organs are present in female flowers, while a pronounced central dome in male flowers may be a remnant of the gynoecium (*39*). Overall, our results tend to support the idea that the most recent ancestor of all extant angiosperms was indeed bisexual.

## Supporting information

Datas2

## Acknowledgments

we thank the Environmental Service of the Northern Province of New Caledonia for permission for seed collection and the staff of the IAC for horticultural assistance, John Bowers for performing some of the earlier work in genome assembly on the *Amborella trichopoda* genome used in this study, and Elise Lucotte for discussions.

## Author contributions

Conceptualization: GABM, CS, JK, JLM, AB Methodology: JK, FP, GABM, JLM, AB Software: JK, FP, Formal analysis: JK, AB, GL, AB, JB, AH Investigation: JK, AB, AAR, GL, JC, CS, GABM, JLM Resources: BF, GG, PR, CWdP Writing - Original Draft: JK, GABM, JLM, CS Writing - Review & Editing: all authors, Visualization: JK, AB, Supervision: GABM, JLM, CS, JK Project administration: GABM, JLM, CS, Funding acquisition: GABM, CS, JLM, CWdP ;

## Data and materials availability

The *A. trichopoda* genome is in CoGe (V.6, genome_ID = 50948, https://genomevolution.org/coge), all of the genome re-seq data are in SRA under the bioproject PRJNA212863, the RNA-seq data are in SRA under the bioproject PRJNA687004.

## Supplementary Materials

Materials and Methods

Supplementary

Text Figures S1-S3

Tables S1-S2

