## Supplementary material file for "A derived ZW chromosome system in *Amborella trichopoda*, the sister species to all other extant flowering plants"

**This PDF file includes:**

Materials and Methods
Supplementary Text
Figs. S1 to S3
Table S1 to S2

**Other Supplementary Materials for this manuscript include the following:**

Data S1: excel file including all the genes in the SDR region and information about them
Data S2: excel file including annotation for all the chromosome 9 genes

### Materials and Methods

Plant material

We used RNA-seq data for 10 female and 10 male individuals grown from seeds collected in the wild in New Caledonia (Mont Aoupinié). Seeds were sown in New Caledonia, and after germination, plants were sent to Lyon (France) to be cultivated in the greenhouse facilities of the ENS Lyon.

16 female and 17 male F1 siblings were generated from mating a single female and male individual (courtesy of Ron Determann at the Atlanta Botanical Gardens, Atlanta GA, USA).

Sequencing

RNA was extracted from several flower buds of each plant with the Spectrum (TM) Plant Total RNA Extraction kit (Sigma-Aldrich), and sent to the Genoscope for sequencing on a Solexa platform (100 bp, paired-end).

Whole genome resequencing using Illumina Paired-End (PE) 150 base pair (bp) was generated for each F1 individual.

Read processing and assembly

RNA-seq reads were processed through removal of adapter sequences, quality filtering (>20), and filtering of ribosomal sequences.

SNP calling and *SDpop* analysis

RNA-seq reads were mapped onto the annotated V6.1 draft genome using STAR (version 020201). Only reads mapping to annotated genes were retained in further analysis, and individuals were genotyped using these reads with reads2snp (version 2), allowing for an allelic expression bias, and without using cleaning for possible paralogs.

From these genotypes, the sex-determining system was inferred using the novel method SD-Pop (*24*). In short, assuming panmixia, for each SNP, we assessed the most likely equilibrium between allele and genotype frequencies: 1) autosomal segregation modelled as Hardy-Weinberg equilibrium for diploidy; 2) haploid segregation for which the allele frequency equals the genotype frequency; 3) paralogous segregation modelled as Hardy-Weinberg equilibrium for tetraploidy; 4) sex chromosome hemizygosity modelled as Hardy-Weinberg equilibrium for the homogametic sex and haploid segregation for the heterogametic sex; and 5) sex chromosome allosomy modelled as modified Hardy-Weinberg equilibria taking different allele frequencies between the homologous, non-recombining chromosomes into account. The proportion of sites in the genome for these five segregation types, as well as a genotyping error rate, were optimized using an Expectation-Maximization (EM) algorithm. The probabilistic framework of this method yields the likelihood of the data for the absence of sex chromosomes, or for an XY or a ZW system; these likelihoods are compared using the Bayesian Information Criterion (BIC). Furthermore, for each SNP, the posterior probabilities associated to each segregation type are calculated, as well as for each contig in the annotation.

SNP calling, SNP density and heterozygous site analyses

Re-seq read were mapped to the *A. trichopoda* v6.1 genome assembly using bwa v0.7.15 (mem) with default parameters. Single Nucleotide Polymorphisms (SNPs) were identified within the F1 mapping population using freebayes-parallel v1.2.0 (parameters: -C 5 -3 300 -p 2 -m 30 -q 30 -n 4).

Nucleotide diversity measured as *pi* per-site was estimated using VCFtools v.0.1.15 among female and male F1 siblings and an average and 95% Confidence Interval (CI) was estimated every 50 kilobase pairs (Kbp) with 25 kb sliding across the entire *A. trichopoda* v6.1 genome assembly.

Similarly, heterozygosity among female and male F1 siblings was estimated using VCFtools v.0.1.15. Comparison of average *pi* per-site and 95% CI per bin (and heterozygosity) between female and male F1 siblings was used to identify diverged genomic regions between sexes. Specifically, high levels of *pi* per-site or heterozygosity in among females compared to males is indicates the presence of a ZW sex chromosome system (female heterogameity), whereas if the opposite is observed, indicates the presence of a XY sex chromosome system (male heterogameity).

Male/female coverage analysis

Whole genome resequencing from the F1 mapping population was used to estimated female and male average sequencing coverage and 95% CI every 20 Kbp with 10 Kbp sliding across the entire A. trichopoda v6.1 genome assembly.

Putative W-specific regions were identified as F1 female average sequencing coverage being $\geq\log_2(2)$ and approximately statistically significant (re. lower bound 95% CI being greater than F1 male upper 95% CI) than F1 male average and 95% CI sequencing coverage. Nested and/or flanking regions in/next to W-specific regions that were approximately statistically significant but did not meet the $\geq\log_2(2)$ coverage threshold were incorporated into the putative W-specific regions. For contigs that were not incorporated into the large 13 pseudo-molecules (chromosomes), average coverage in F1 female had to be ≥1 before the described logic to identify W-specific regions was applied.

Putative shared Z regions were identified as F1 male average sequencing coverage being $\geq\log_2(1)$ and is approximately statistically significant (re. male lower bound 95% CI is greater than female upper 95% CI) than F1 female average and 95% CI sequencing coverage. The same logic was applied to nested and/or flanking regions. However, for contigs, average coverage of both males and females had to be ≥1 before the logic was applied.

Gene content of the sex-determining region

InterProScan v5.32-71.0 (parameters: -appl TIGRFAM,PANTHER,SMART,CDD,Pfam -f TSV -goterms -iprlookup) was used to identify protein functional domains and Gene Ontology (GO) terms for each annotated protein coding gene found within and outside of the W-specific and shared Z regions. Biological Process (BP), Cellular Component (CC), and Molecular Function (MF) GO enrichment compared to GO terms associated with all annotated protein coding genes for W-specific and shared Z regions annotated protein coding genes was estimated using a weighted Fisher's exact test ($P < 0.05$) in the R package topGO.

Unique Pfam protein functional domains were identified between W-specific and shared Z regions annotated protein coding genes. Additionally, abundance of unique Pfam protein functional domains was estimated between W-specific and shared Z regions annotated protein coding genes.

*Arabidopsis thaliana* homologs to the W-specific and shared Z regions annotated protein coding genes were identified using BLASTp (parameters: -evalue 1E-08 -max_hsps 1 -

max_target_seqs 1). Described *A. thaliana* phenotype(s) for each best BLASTp hit to W-specific and shared Z regions annotated protein coding genes was extracted from The Arabidopsis Information Resource (TAIR) (https://www.arabidopsis.org/).

Synonymous divergence analysis

The existence of both Z and W copies of homologous regions in the assembly makes it difficult to compare Z and W copies directly. Thus, the RNA-seq and resequencing data were mapped on the assembly with either the putative W-specific regions or the putative Z-specific regions (identified as described above) masked, using otherwise the same mapping and genotyping protocols as mentioned before (STAR and reads2snp for RNA-seq data, bwa and freebayes for the mapping population resequencing data). For sex-linked SNPs, Z and W alleles were identified from SDpop's and SEX-DETector's output. These allow, for the population sample (RNA-seq data) to calculate the divergence between Z and W, and the nucleotide diversity of each copy. For both datasets, Z and W haplotypes were reconstructed from the sex-linked SNPs; in the RNA-seq data, for alleles that were not completely fixed, the allele that exceeded a frequency of 0.6 was incorporated in the corresponding haplotype. Then, codeML was run in pairwise mode on each Z-W haplotype pair to calculate dN, dS and dN/dS.

**Supplementary Text**

Weak association to sex on the left arm of chromosome 9

In region 10-20 Mb, males tend to be heterozygous and females homozygous in a pattern reminiscent of *XY* systems, though sex-association was much weaker than observed for the *ZW* region (Figure 1). The seed parent of the mapping population exhibited higher heterozygosity in this region than the pollen parent, but counter to expectations F1 offspring exhibited male-biased heterozygosity (Figure S1). In our mapping population, we hypothesize elevated recombination between the sex-determining region of chromosome 9 (45-49 Mb) and the pseudo-autosomal regions to be the primary mechanism for male-biased heterozygosity in the region to the left of the pericentromeric region (10-20 Mb) (Figure S1). Re-sequencing analysis of six progeny sired by a second male with lower heterozygosity in this region revealed female-biased heterozygosity consistent with this possible explanation (Figure S1).

Resolving W and Z chimerism in the sex-determining region of the A. trichopoda v6 genome

Mapping of the DNA resequencing read to the published genome revealed important variation in the male and female coverage in the non-recombining region. More precisely, some regions had (almost) no reads from males, while in others, about two thirds of the reads were from males. This pattern is expected for W-specific and Z-hemizygous genes, respectively. However, several genes with no male reads were very similar to genes for which two thirds of the reads were from males, which could be due to the inclusion of both the Z and the W gametolog in the reference assembly. To solve this issue, we produced two masks of the assembly, based on coverage differences between females and males (accompanying figure). In one mask of the assembly, parts with only female coverage are removed (the "NoW assembly"), and in the other, parts with a two-fold increase in male coverage compared to females were removed (the "NoZ assembly").

The masked assemblies ensure that reads from both Z and W copies, if they exist, are mapped on the same reference sequence, thus allowing to quantify divergence between the haplotypes.

Remapping the DNAseq reads to these assemblies removed most variation in the coverage (accompanying figure). However, a few genes still had substantially sex-biased coverage, either male-biased (two thirds of reads come from males) or female-biased (no male reads). These could be W-specific or Z-hemizygous genes (Table S2), although some of these might also be caused by Z and W gametologs that were present in the genome after the masking.

**Fig. S1.**

Patterns of transmission of heterozygosity from parents to offspring and interpretation of the data assuming a hot spot of recombination between region 10-20 Mb and the SDR in family Am_F3 x Am_3_2 i.e. the mapping population mentioned in the main text (A) and another family with the same mother and a different father Am_F3 x Am_1_2 (B). A) The offspring female – male heterozygosity shows that the heterozygosity in the mother is observed in the sons (green bars). This can be explained if a strong hot recombination hotspot between region 10-20 Mb and the SDR (region 45-49 Mb) resulted in the production of recombinant gametes (A,Z and a,W) in the mother. The sons are thus A,Z and a,Z and the daughters a,W and a,Z. B) in the other family the pattern of transmission of heterozygosity from parents to offspring is different. The heterozygosity of the mother is not observed in the sons but in the daughters (green bars). This time, the genotype of the father is different and A,Z / A,Z. The results can be explained again with a hot spot, resulting in recombinant gametes in the mother and heterozygous sites in the daughters.

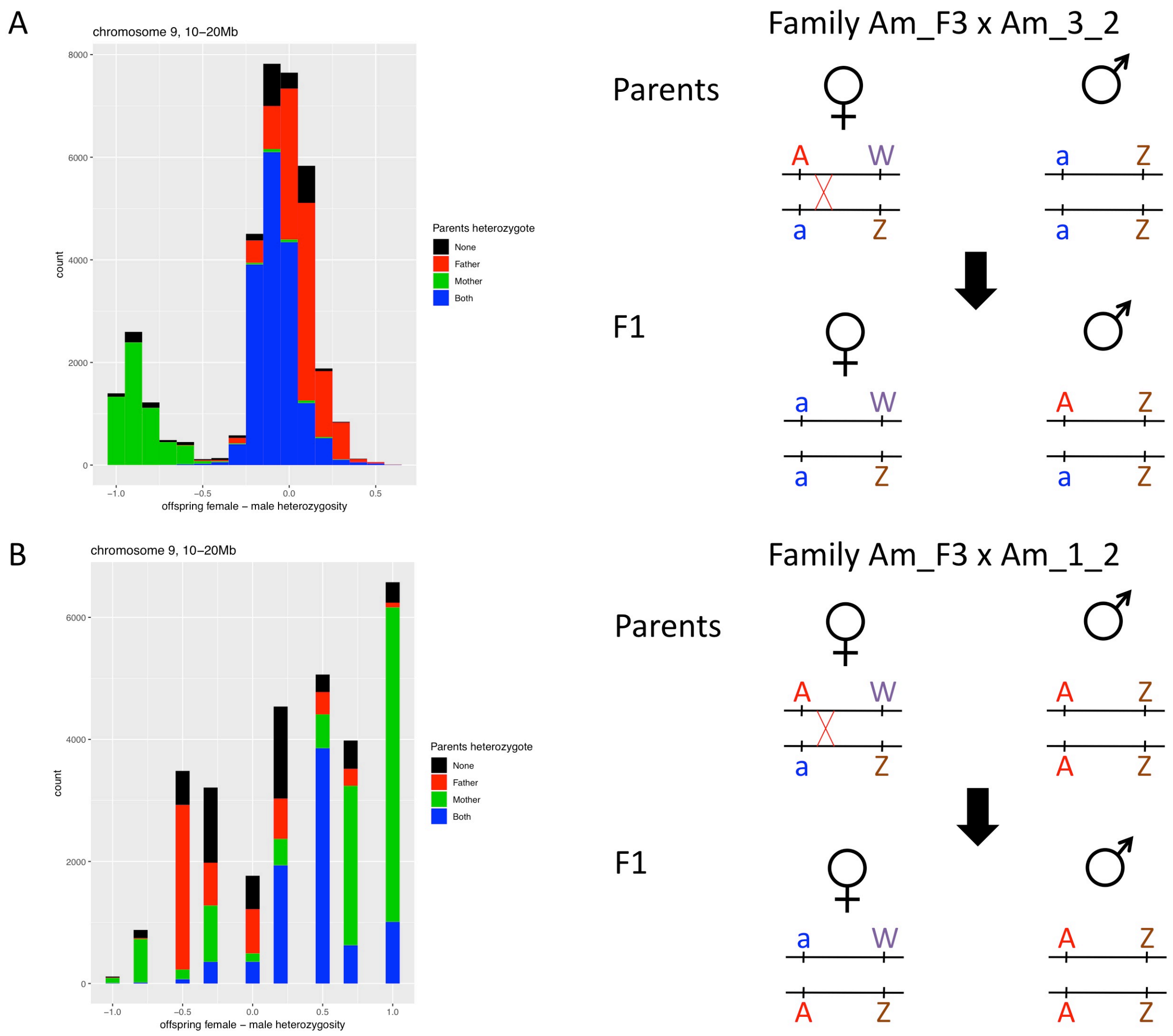

**Fig. S2.**

Average proportion of male reads among the total number of reads per gene in the sex-determining region of chromosome 9. The regions with highly biased read counts when mapping on the raw (unmasked) assembly were used to produce two masked genome assemblies: one ("NoW") with the highly female biased regions removed (indicated in red), and one ("NoZ") with the highly male biased regions removed. The reads were remapped on each of these masked genomes, resulting in much less bias.

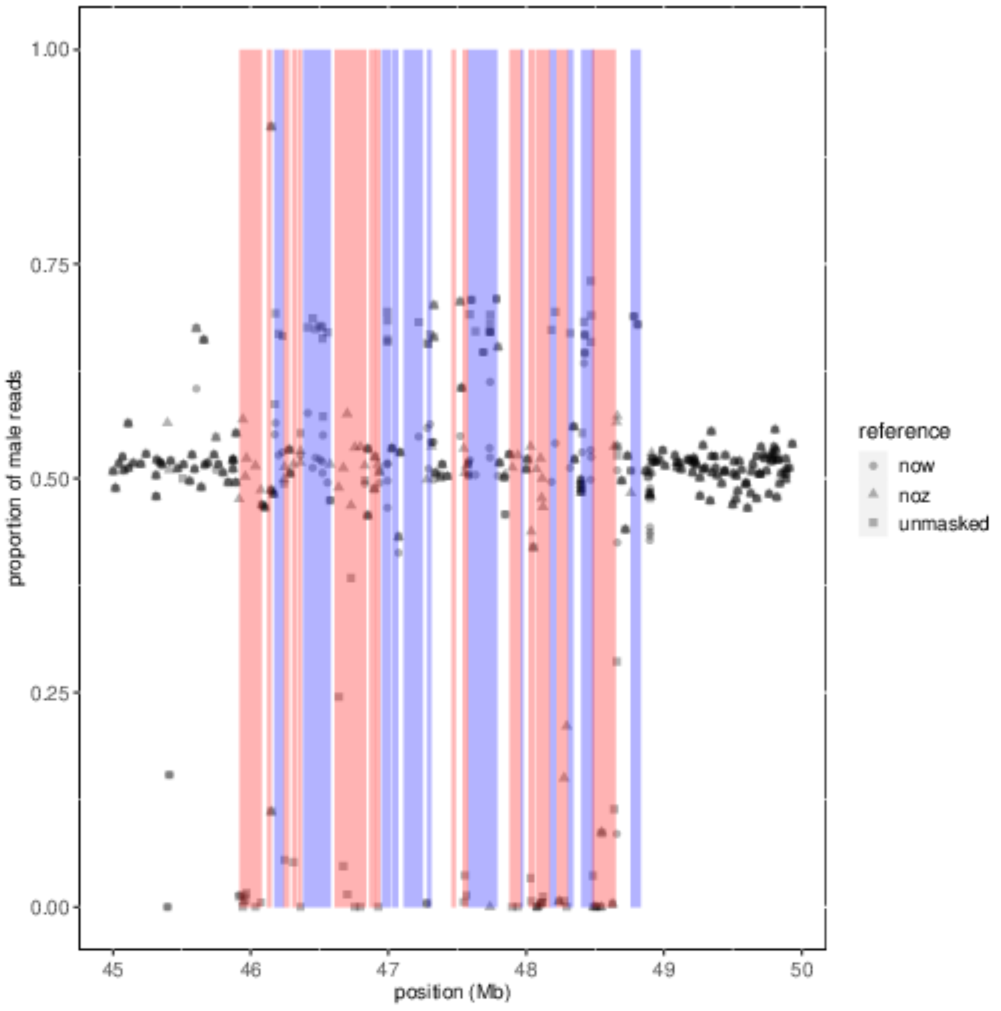

**Fig. S3.**

Sex-biased gene expression in the RNA-seq data of flower buds from a wild population (10 females and 10 males). The fold-change (logFC) was calculated using the male over female expression ratio, thus negative values indicate the females expressed the gene more. P-values were calculated with edgeR.

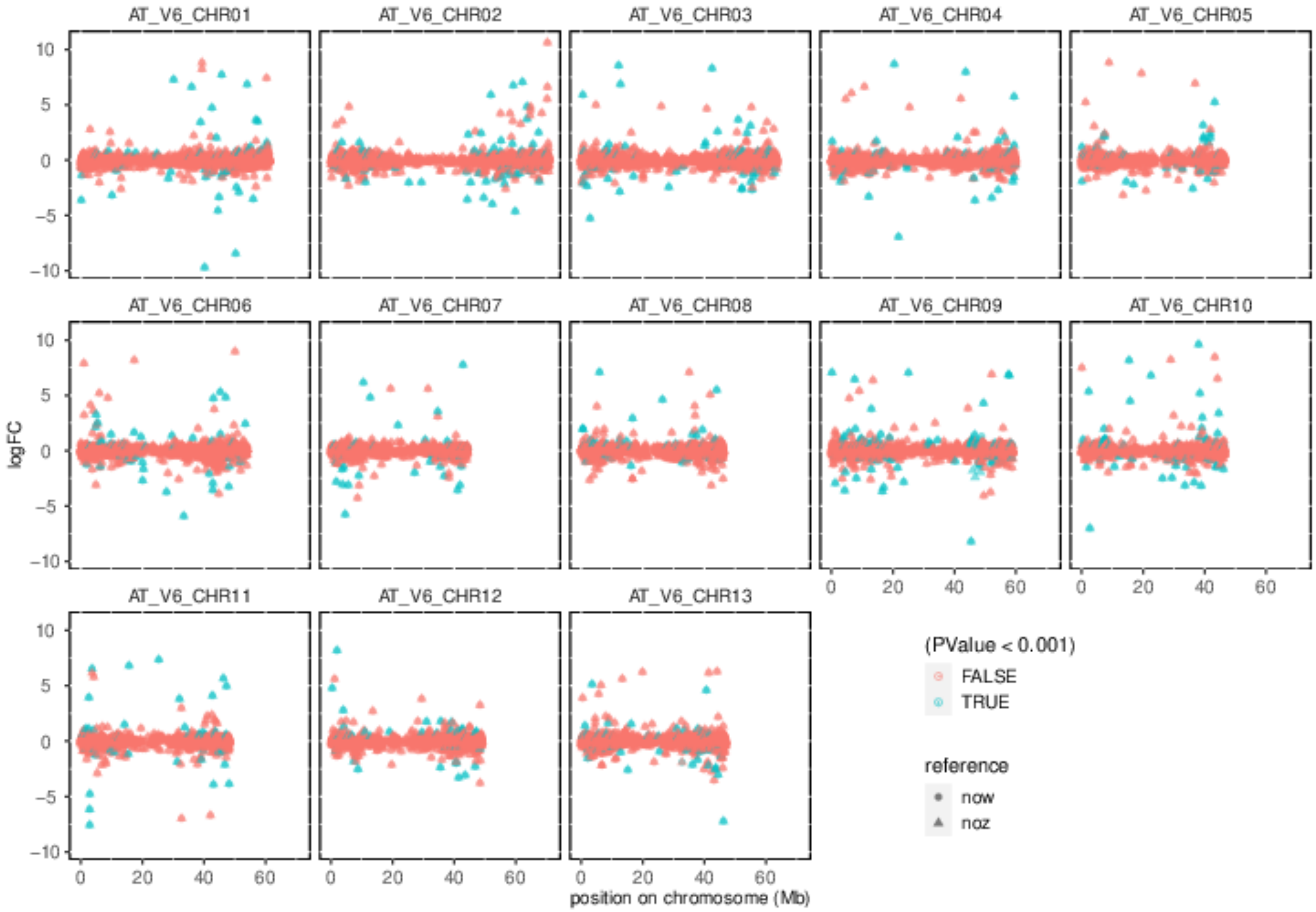

**Table S1.**

Presence of female-specific (putative W) alleles, identified in the RNA-seq data, in the mapping population data. The RNA-seq data, mapped on the unmasked genome (using STAR and genotyping with samtools) yielded 671 alleles for which all (10) males were homozygous and all (10) females heterozygous. These were compared to the alleles at the same genomic positions in the mapping population.

| Names | Sex | Number of W-alleles detected |
|---|---|---|
| Am_3_2 | M (father) | 0 |
| Ambo_005 | M | 0 |
| Ambo_013 | M | 0 |
| Ambo_016 | M | 0 |
| Ambo_017 | M | 0 |
| Ambo_028 | M | 0 |
| Ambo_051 | M | 0 |
| Ambo_054 | M | 0 |
| Ambo_021 | M | 1 |
| Ambo_030 | M | 1 |
| Ambo_035 | M | 1 |
| Ambo_055 | M | 1 |
| Ambo_024 | M | 2 |
| Ambo_026 | M | 2 |
| Ambo_009 | M | 3 |
| Ambo_036 | M | 4 |
| Ambo_022 | M | 5 |
| Ambo_038 | M | 5 |
| Ambo_007 | F | 456 |
| Ambo_014 | F | 465 |
| Ambo_011 | F | 468 |
| Ambo_008 | F | 470 |
| Ambo_006 | F | 482 |
| Am_F3 | F (mother) | 484 |
| Ambo_027 | F | 499 |
| Ambo_010 | F | 504 |
| Ambo_073 | F | 510 |
| Ambo_056 | F | 511 |
| Ambo_068 | F | 512 |
| Ambo_050 | F | 513 |
| Ambo_074 | F | 514 |
| Ambo_015 | F | 519 |
| Ambo_052 | F | 525 |
| Ambo_057 | F | 525 |
| Ambo_071 | F | 526 |

**Table S2.**

List of sex-biased, Z-hemyzygous or W-specific genes located in the sex-determining region with possible role on sex-determination.

| Gene ID | Annotation | Type | Possible effect |
|---|---|---|---|
| AmTr_v6.0_c9.19680.1 | PF14432, DYW domain acid deaminase (PPR gene) | Female-biased, W-specific gene | Feminizing effect |
| AmTr_v6.0_c9.19910.1 | AT2G21220.1 SAUR-like auxin-responsive protein family | Male-biased | Masculinizing effect |
| AmTr_v6.0_c9.20000.1 | AT2G21220.1 SAUR-like auxin-responsive protein family | Male-biased | Masculinizing effect |
| AmTr_v6.0_c9.20030.1 | AT1G75590.1 SAUR-like auxin-responsive protein family | Male-biased | Masculinizing effect |
| AmTr_v6.0_c9.20270.1 | AT1G19850.1 Transcriptional factor B3 family protein / auxin-responsive factor AUX/IAA-related | Male-biased | Masculinizing effect |
